# An Fc-SPINK1 fusion protein inhibits pancreatic inflammation in a mouse model

**DOI:** 10.1101/2025.02.21.639528

**Authors:** J. C. Way, D. Heid, M. Theiss, S. Nørrelykke, L. Barck, L. Burger, J. A. Bredl, T. Dobbertin, P. Höflich, T. Stadelmann, D. Niopek, K. Redfield Chan, R. Sivaraman, A. R. Graveline, A. Vernet, M. Sanchez-Ventura, P. A. Silver

## Abstract

Pancreatitis results from premature activation and impaired inactivation of pancreatic proteases, primarily trypsin, leading to self-digestion, tissue necrosis, fibrosis, and inflammation. SPINK1 is a pancreas-specific inhibitor of trypsin that prevents premature trypsin activation, and could be a candidate therapeutic. However, because of its small size, SPINK1 would be subject to rapid renal clearance, making it ineffective. To construct a long half-life therapeutic inhibitor of trypsin for pancreatitis treatment we fused this protein to the C-terminus of an IgG1 antibody Fc element, increasing the size to ∼78 kDa, thereby exceeding the renal clearance threshold and providing for FcRn-mediated recycling out of cells. A non-glycosylated form of Fc-SPINK1 was expressed in the yeast *Pichia pastoris*. Fc-SPINK1 inhibits trypsin enzyme activity *in vitro*. The blood pharmacokinetics in mice are consistent with a three-compartment distribution model and a terminal half-life of ∼3 days. In a caerulein-induced mouse model of pancreatitis, Fc-SPINK1 significantly ameliorated cell death and immune cell infiltration. We developed an automated image analysis technique to quantify pancreatitis-associated loss of tissue cohesion, and found that Fc-SPINK1 also reduced this effect. This study demonstrates the potential of Fc-SPINK1 as a rationally designed therapy for pancreatitis.

## Introduction

Pancreatitis is a major unmet medical challenge, affecting nearly 9 million people annually and resulting in more than 100,000 deaths/year.^[1–2]^ Acute pancreatitis has a high mortality, in the range of 3-16%.^[3–6]^ Patients with chronic pancreatitis can have acute attacks, can also progress to pancreatic cancer, and may also have inflammatory diabetes that is often mistaken for Type II diabetes. The disease is influenced by genetic factors, alcohol consumption, trauma, gallstones and also occurs as a complication of endoretrocholiangopancreatographic surgery (ERCP).^[7]^ Gallstone-driven pancreatitis is well-treated by surgery, and acute pancreatitis involves a SIRS-like cytokine storm that is challenging to treat in any case. In contrast, chronic and ERCP-induced pancreatitis have a predictable course that may allow for pharmacological intervention.

Genetic studies have demonstrated the central role of trypsin in the development of pancreatitis, and point to a multicomponent regulatory system for controlling trypsin activity.^[8]^ Trypsin cleaves the pro-form of other digestive enzymes including prochymotrypsin. Mature chymotrypsin can then cleave and inactivate trypsin. In addition, the small, pancreas-specific protein SPINK1 inhibits trypsin. Gain of function mutations in cationic trypsinogen (PRSS-1) and loss of function mutations in SPINK1 and chymotrypsin significantly increase the risk of pancreatitis. ^[9–12]^ In the pancreatic acinar cells, pro-trypsin, pro-chymotrypsin, other digestive enzymes, and SPINK1 are co-localized in vesicles that normally fuse with the apical surface to release their contents into a duct system that leads to the small intestine. In pancreatitis, these vesicles aberrantly fuse with lysosomes, and the lysosomal hydrolase cathepsin B activates trypsin, initiating a chain reaction and autolysis of acinar cells with the destruction of surrounding tissue. ^[13–15]^ Studies in mice associate the loss of cathepsin B with reduced trypsin activity and milder pancreatitis.^[13][14]^

Various agents have been investigated for the treatment of pancreatitis, including antisecretory molecules, protease inhibitors, calcium modulators, immunomodulators, anti-inflammatories, and antioxidants. For trypsin inhibitors, the small protein Aprotinin/BPTI and small molecule gabexate mesylate were both extensively clinically tested against acute pancreatitis and generally failed,^[16–18]^ most likely because both of these molecules have a very short plasma half-life (2 hours and <1 minute, respectively),^[17–19]^ and because in acute pancreatitis the risk of death is likely driven by inflammatory cytokines, and it is too late to address the initiating problem of trypsin overactivity.^[20]^ However, clinical trials have shown that 13-hour or 6.5 hour infusions of gabexate mesylate prior to ERCP did significantly reduce the incidence of pain and acute pancreatitis after the procedure.^[21–22]^ Unfortunately, the long infusion times before surgery make this specific treatment impractical. These trials demonstrate that trypsin inhibition could be of use in prophylaxis of pancreatitis before ERCP, but also underscore the importance of pharmacokinetics in driving adoption of a particular treatment.

As the human body’s natural endogenous trypsin inhibitor, SPINK1 (serine protease inhibitor Kazal type 1) is a promising therapeutic element. Compared to BPTI, SPINK1 has the advantage that it is an endogenous human protein and less likely to induce an immune response after repeated administration in treatment of chronic disease. In this work, we demonstrate that an Fc-SPINK1 fusion protein can reduce the severity of pancreatitis in a mouse model.

## Results

### Design, production, and purification of Fc-SPINK1, and *in vitro* trypsin inhibition

The trypsin inhibitors SPINK1 and BPTI (bovine pancreatic trypsin inhibitor; Aprotinin) both have a molecular weight of <7,000 Daltons, which will result in rapid renal clearance and to make these proteins unsuitable as drugs. We fused SPINK1 and BPTI to the C-terminus of the Fc fragment of human IgG to create an Fc-SPINK1 and Fc-BPTI dimers with a molecular weight of about 78 kDa, which is well above the renal clearance threshold and should significantly improve pharmacokinetics^[23]^ (Figure 1a-c).

**Figure 1:**
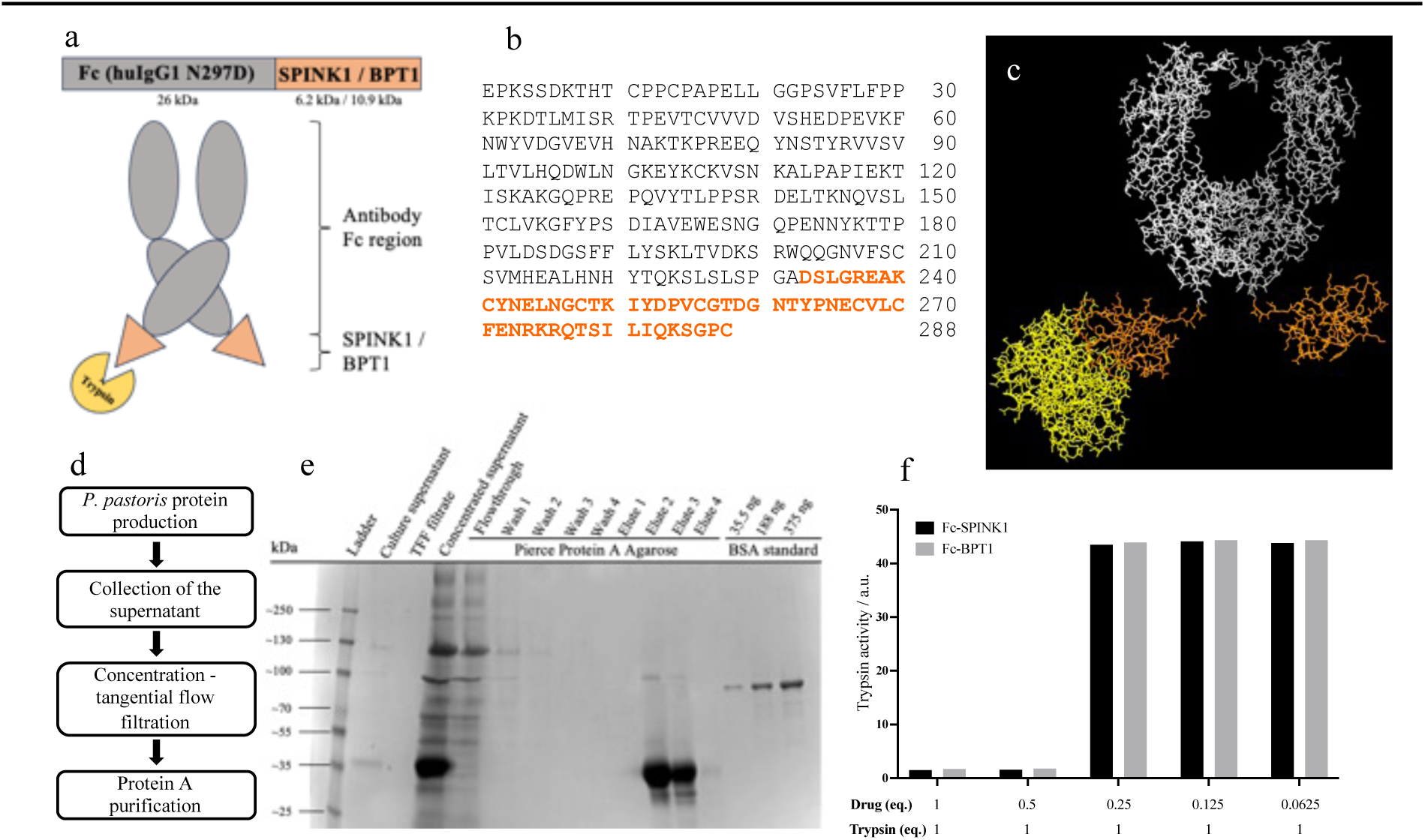
The fusion proteins Fc-SPINK1 and Fc-BPTI are constructed from the Fc region of a human IgG1 antibody and SPINK1 resp. BPTI. SPINK1 is a naturally derived human trypsin inhibitor, BPTI is found in bovine pancreas. (a) Schematic structure of Fc-SPINK1 or Fc-BPTI. (b) Sequence of the mature, secreted form of Fc-SPINK1. (c) The predicted 3D structure of Fc-SPINK1 and Fc-BPTI. (d) Overview of the production and purification procedure of Fc-SPINK1 and Fc-BPTI in the yeast *Pichia pastoris*. (e) The major production and purification steps are visualized using Coomassie stained SDS-PAGE. The data displays that Fc-SPINK1 (32 kDa) was effectively purified and concentrated. (f) The trypsin inhibition assay demonstrates the inhibition of trypsin by Fc-SPINK1 and Fc-BPTI in a para-nitrophenol-based assay. Both constructs successfully inhibit trypsin at a 1:1 and 1:0.5 ratio.

The Fc region contains three mutations relative to the natural human IgG1 sequence (Figure 1b). First, amino acid 5, normally cysteine, is mutated to serine because this cysteine normally forms a disulfide bond with the antibody light chain, which is not present in this construct. Second, the aspartic acid at position 82 is normally asparagine, but this is the site of N-linked glycosylation of the Fc region. Removing this glycosylation has the effect of significantly reducing binding to both Fc receptors and to the complement component C1q,^[24]^ and also allows for safe, low-cost expression in the yeast *Pichia pastoris* instead of mammalian cells; specifically, *Pichia* generates a high-mannose-type N-linked oligosaccharide modification that would be recognized as foreign by mammals,^[25]^ but our molecule has been engineered to not be glycosylated. Third, amino acid 232 is normally the C-terminal lysine of a secreted IgG1, but this amino acid can be cleaved in both natural antibodies and fusion proteins, so it is mutated to alanine to prevent potential cleavage.^[26]^ The SPINK1 and BPTI modules were placed at the C-terminus of Fc instead of the N-terminus, because fusions in the Fc-X orientation are generally expressed at a higher level.^[23]^ No linker was used because the C-terminus of Fc is somewhat unstructured and may form a natural linker, and because both fusion termini are far from known sites of protein interaction (Figure 1c).

The *Pichia pastoris* expression system was used for the production procedures, and fusion proteins were purified to near homogeneity by protein A chromatography (Figure 1d-e; Methods). A trypsin inhibition assay (Figure 1f) demonstrated the efficacy of both constructs in inhibiting trypsin at molar ratios of 1:1 (0.429 nM each) and 1:0.5 Fc-SPINK1-trypsin. (We generally did not observe partial inhibition of trypsin by Fc-SPINK1 at low molar ratios of SPINK1:trypsin; we hypothesize that trypsin might inactivate Fc-SPINK1 under the conditions we used.)

### *In vivo* efficacy in caerulein-induced mouse model of acute pancreatitis

Fc-SPINK1 and Fc-BPTI were evaluated as potential therapeutic agents in a mouse model of caerulein-induced pancreatitis. Caerulein is a frog-derived toxic peptide that stimulates a moderate, transient pancreatitis.^[27–28]^ Pancreatitis was induced by seven injections of caerulein into C57Bl/6N mice, with a single injection of Fc-SPINK1, Fc-BPTI, or vehicle, followed by harvest of the pancreas and blood 18 hours after the first injection (Fig. 2A). Pancreases were fixed, sectioned, and stained with hematoxylin/eosin (H&E) or with anti-CD11b (staining macrophages, neutrophils, and certain other immune cells^[29]^) (Fig. 2B-G). To date, necrosis and immune cell infiltration is quantified visually. This slow process requires subsampling of tissue.^[41]^ To increase statistical power, we performed semi-automated analysis on all available image slides with respect to necrotic cells (Fig. 2H-I), and immune cell infiltration (Fig. 2J-K). In H&E-stained healthy pancreas tissue, the acinar cells are normally tightly associated with each other (Figure 2B). After cerulein treatment, the cells are dramatically separated in H&E-stained sections (Figure 2C).^[30]^ To quantitate this effect in an unbiased way, we developed a new automated image analysis method (Fig. 2L-M; see below).

**Figure 2.**
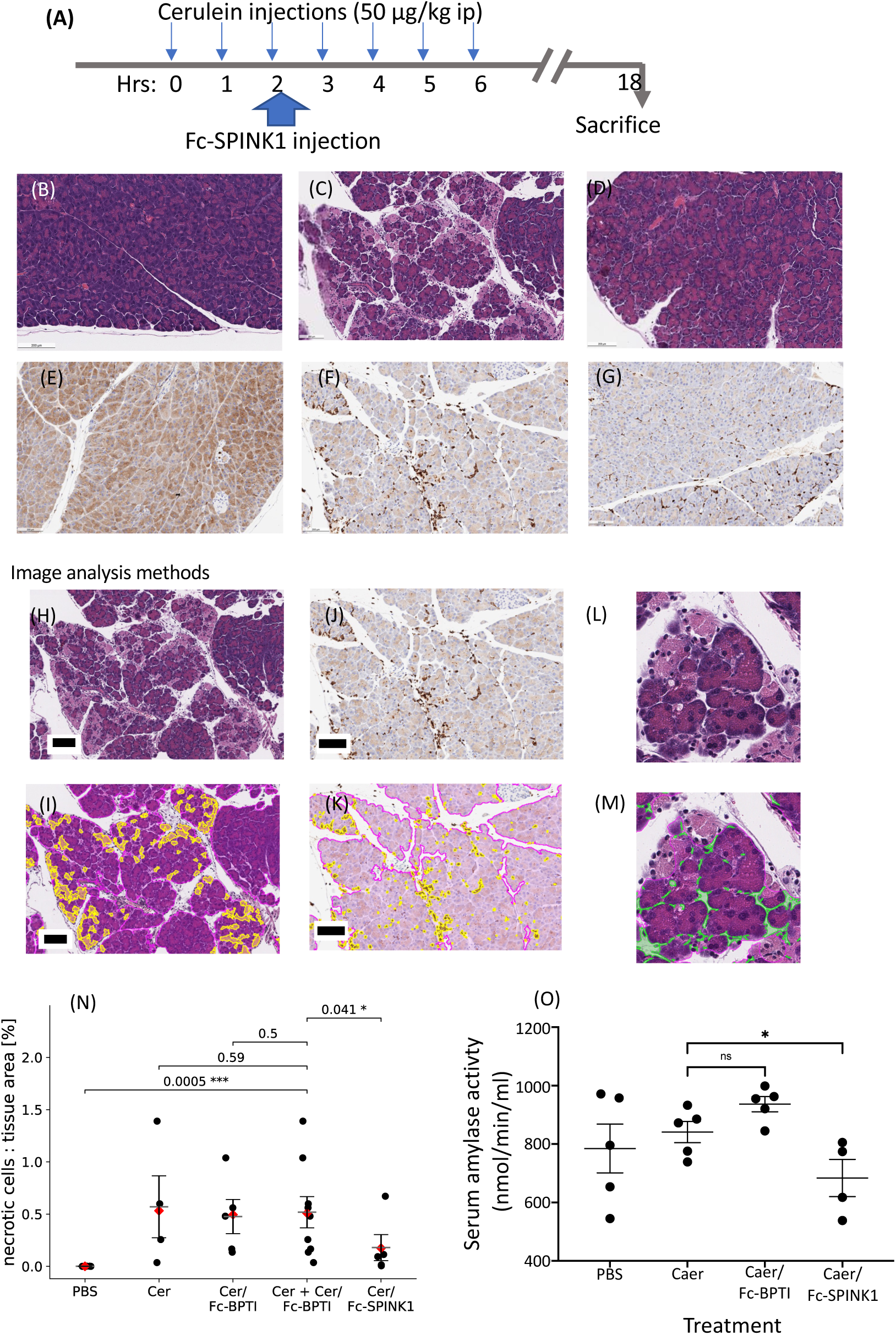
Treatment of cerulein-induced pancreatitis with Fc-SPINK1 reduces symptom severity. **(A)** Workflow of the experiment. N=5 C57Bl6N mice per dose group. **(B-D)** Hematoxylin/Eosin staining of pancreas tissue sections from mice treated with **(A)** seven hourly injections of PBS, **(B)** seven hourly injections of ceruelin, or **(C)** seven hourly injections of ceruelin plus an injection of 5 mgs of Fc-SPINK1. The results indicate that in cerulein-treated mice, a number of cells beome necrotic (light pink staining), and cells also suffer from a loss of junctional integrity, going from a cuboid shape to a rounded shape with fluid between cells. Treatment with Fc-SPINK1 may lessen these phenotypes. **(E-F)** Anti-CD11b staining of pancreas tissue sections from mice treated with PBS, cerulein, or cerulein plus Fc-SPINK1. The results indicate that in cerulein-treated mice, CD11b+ leukocytes (e.g. macrophages and NK cells) invade the pancreas. Treatment with Fc-SPINK1 may lessen this phenotype. **(H-M)** Image processing methods for automated, unbiased evaluation of stained tissue sections. See Supplementary Information for algorithmic details. Scale bars are 100 microns. **(H-I)** Semi-automated scoring of cell necrosis. Yellow false-colored areas are those scored as necrotic. Magenta false-colored area indicates exocrine tissue **(J-K)** Automated scoring of CD11b+ infiltration. Yellow false-colored cells are CD11b+. Magenta false-colored cells indicate exocrine tissue. **(L-M)** Automated scoring of loss of junctional integrity in the pancreas. This algorithm segments gaps between cells to generate a network pattern (green false-color) and then calculates the number of branch points in the network (normalized to the area of the tissue section; Figure 5). **(N)** Results of semi-automated necrotic cell staining. Pancreases of PBS-treated mice show no necrotic cells, while pancreases of cerulein treated mice showed a increased level of cell necrosis. Injection of Fc-SPINK1 appeared to lessen cell necrosis, while Fc-BPTI had no apparent effect on cell necrosis. To improve the statistical analysis, the scores of mice treated with cerulein alone and cerulein plus Fc-BPTI were combined and compared to scores from mice treated with cerulein plus Fc-SPINK1. Horizontal bars indicate mean ± standard error of mean (SEM). Red diamonds indicate mean weighted by tissue-area. P-values: one-tailed Mann-Whitney-U test. **(O)** Plasma amylase levels in mice treated with PBS, cerulein, and Fc-BPTI or FC-SPINK1, 18 hours after the first injection of cerulein. Levels of amylase in mice treated with cerulein and Fc-SPINK1 were lower than in mice treated with either cerulein or cerulein and Fc-BPTI (*p<0.05, one-tailed Student’s T-test).

In one experiment, we treated mice with 5 mg of Fc fusion protein after the third caerulein injection. This high dose was based on the fact that mammals synthesize large amounts of trypsin, and only a small amount of the injected Fc-SPINK1 fusion protein will distribute into the pancreas (see Discussion). Fc-SPINK1 reduced markers of caerulein-induced pancreatitis (Figure 2). Specifically, acinar cell necrosis and plasma amylase levels were reduced in cerulein+Fc-SPINK1-treated mice compared to mice treated with caerulein and vehicle or Fc-BPTI (Figure 2N-O). Quantitative semi-automated image analysis of necrotic cells showed a significant reduction in the Fc-SPINK1-treated group compared to the caerulein control (Figure 2, Supplementary 1). Fc-BPTI did not appear to reduce acinar cell necrosis or amylase levels.

Pancreatic amylase, which is released during caerulein-induced inflammation, is a human biomarker for the detection of pancreatic injury. Blood plasma amylase activity (Figure 2D) was reduced in the Fc-SPINK1-treated group (*p < 0.05). There was no reduction in amylase activity in the plasma of Fc-BPTI-treated mice. Typically, in the caerulein-induced pancreatitis mouse model, plasma amylase levels peak after the last caerulein injection.^[30]^ In this experiment, at the 18 hour time-point, the amylase levels are returning to baseline, but the effect of Fc-SPINK1 can still be observed.

In a second experiment, the effects of different Fc-SPINK1 doses administered by intravenous (IV) and intraperitoneal (IP) injection were investigated (Figure 3 and Supplementary Figure 2), using doses of 1, 2.5, or 5 mg per 20-gram mouse. A dose response was observed with the IV-injected mice. Mice receiving 2.5 or 5 mg/mouse showed significantly less necrosis than mice treated with 0 or 1 mg, and the tissue showed progressively ameliorated junctional integrity with increasing IV dosing. In the intraperitoneally dosed animals, the separate dose groups did not show significantly less cell necrosis than the untreated group, but there was a trend towards a therapeutic effect (Supplementary Figure 2). Taken together, the results indicate that pancreatitis symptoms decrease with high enough doses of Fc-SPINK1, and that a maximally effective dose may not have been reached.

**Figure 3:**
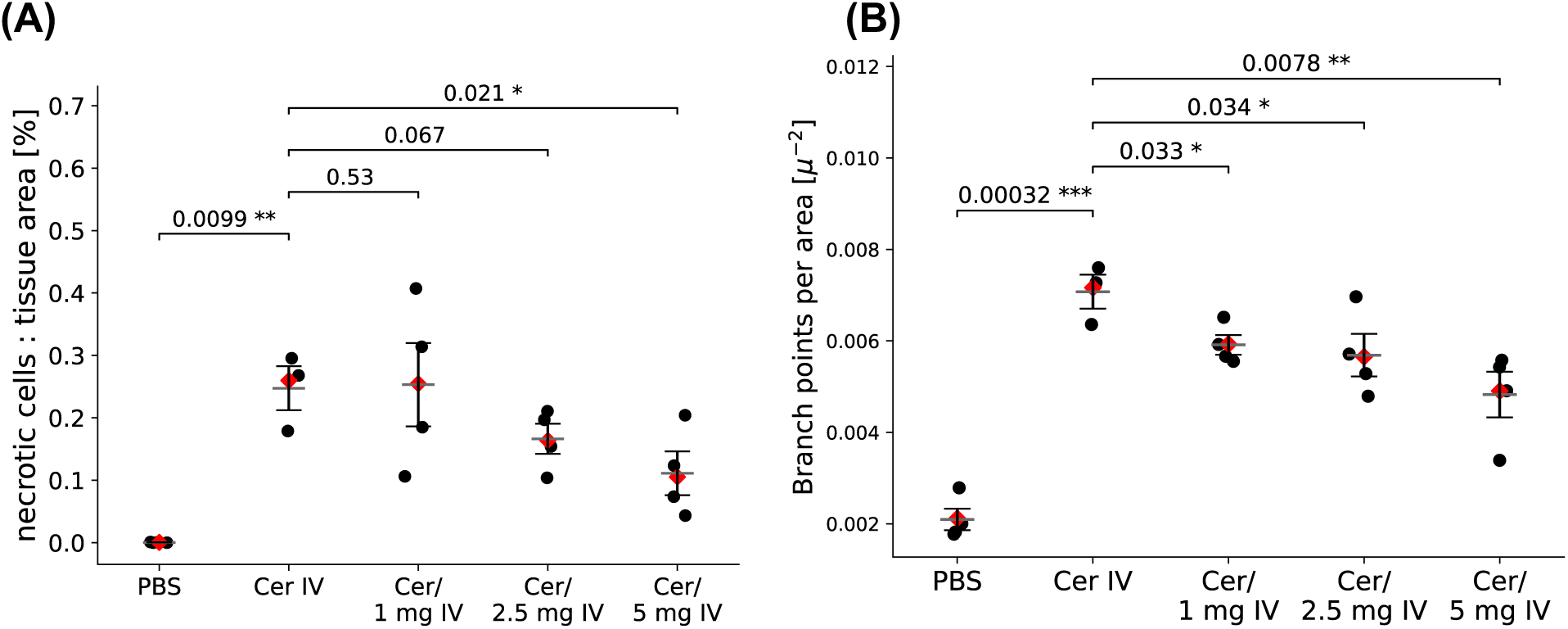
Dose-responsive reduction in cerulein-induced pancreatitic necrosis and loss of tissue integrity. C57Bl6N mice were treated with seven injections of cerulein and one injection of Fc-SPINK1 as in Figure 2A. Horizontal bars indicate mean ± SEM. Red diamonds indicate mean weighted by tissue-area. P-values: One-tailed Welch’s t-test.

Mice of the C57Bl6J strain develop a more severe cerulein-induced pancreatitis than C57Bl6N mice. In a third experiment, when C57Bl6J mice were treated with cerulein and Fc-SPINK1 following the timeline in Figure 2A, a moderate reduction in disease severity was observed. However, when the C57Bl6J mice were given Fc-SPINK1 by IP injection prior to the caerulein injections, we observed a stronger effect, seeing both a statistically significant reduction in cell necrosis and cell separation in the pancreases of caerulein-treated mice (Figure 4). Prophylactic treatment is particularly relevant to ERCP-induced pancreatitis.

**Figure 4.**
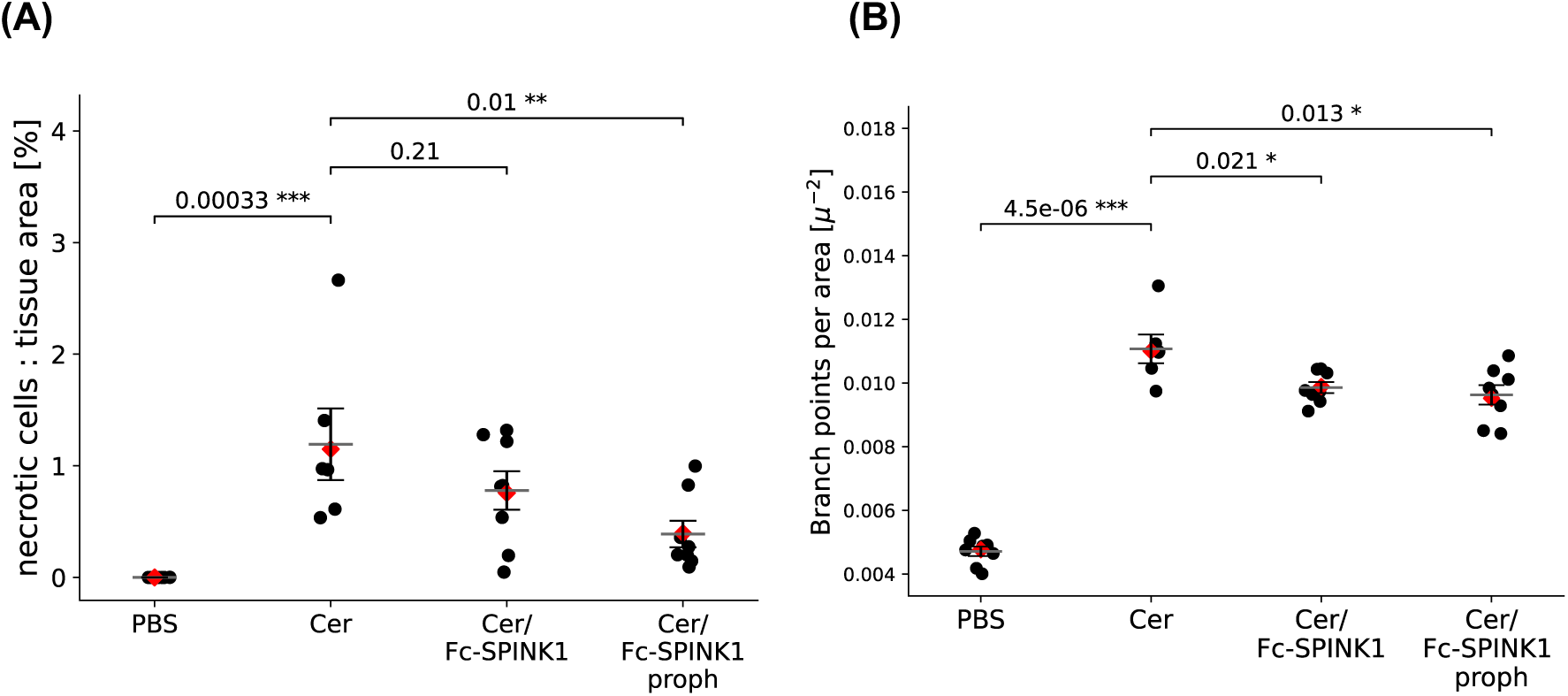
Prophylactic treatment with Fc-SPINK1 reduces effects of cerulein-induced pancreatitis. For both panels,* is p<0.05; ** is p<0.01; ** is p<0.001. **(A)** Necrotic cells were quantitated in H&E-stained pancreas cross-sections, in which mice were treated with Fc-SPINK1 either during the series of cerulein injections as in Figure 2A, or before the first cerulein injection (“prophylaxis”). Black dots represent necrosis scores, and red diamonds represent the mean score, weighted by sample area. P-values: One-tailed Mann-Whitney-U test. **(B)** Branchpoint analysis was applied to the same images. Black dots represent branchpoint scores, and red diamonds represent the mean score weighted by the area of each cross-section. The results suggest that prophylactic treatment with Fc-SPINK1 has a stronger effect on inhibition of cerulein-induced pancreatitis than treatment after initial cerulein injections. P-values: One-tailed Welch’s t-test.

### Image analysis

We used automated and semi-automated image analysis to quantitate the extent of cell death, lymphocyte infiltration, and cell separation, which are visual markers of inflammation. Cell separation may be related to edematous fluid influx, and is henceforth also referred to as edema.

Each analyzed image corresponds to a whole-pancreas cross-section. Images that showed staining-defects or were acquired with different microscope-settings than the rest of their batch, were excluded from analysis. First, exocrine tissue (acinar cells) was segmented using a pixel classifier in QuPath.^[42]^ A random-forest based pixel classifier assigns a class to each pixel (such as background, exocrine tissue) based on rules learned from sparse manual training annotations. Manual corrections were made by removing wrongly segmented (false positive) areas. To ensure pixel classifiers for Edema and Necrosis were trained on well-randomized samples, square regions at a random location within exocrine tissue were automatically selected and combined into a training image.

The Edema, Necrosis, and CD11b+ scoring functions were generated as follows (see Supplementary Information for details). Within exocrine segments, the space between cells was segmented using a QuPath pixel classifier. In inflamed tissue, acinar cells are rounded, possibly due to a loss of tight junctional integrity that may relate to edematous fluid influx into tissue or to separation of cells during fixation. Hence, the intercellular space forms a netlike topology. Branch points signify a loss of tight junction integrity at multicellular junctions. For the Edema function, the number of branch points was therefore used to quantify inflammation. For the Necrosis function, necrotic cells were segmented using a QuPath pixel classifier, and false positive segments were manually removed. The ratio of necrotic cells to exocrine tissue was used to quantify this aspect of pancreatitis. The CD11b+ function is based on the fact that inflammation correlates with CD11B+ immune cell invasion. In IHC images, CD11B+ cells and tissue (exocrine and endocrine) were segmented using a QuPath pixel classifier, and the percentage of tissue area classed as CD11B+ was used to quantify inflammation.

The Shapiro-Wilk test of normality was conducted for each treatment condition to assess the normality of sample distributions, with a significance level of α = 0.05. If the null hypothesis of at least one condition within an experiment was rejected, the Mann-Whitney-U test^[31]^ was used with the alternative hypothesis that sick samples show greater indicators of inflammation than healthy or treated samples. When the null hypothesis of the Shapiro-Wilk test^[32]^ of normality was not rejected, Welch’s one-tailed t-test^[33]^ was applied instead.

### Pharmacokinetics

We investigated the dynamics of Fc-SPINK1 in the blood system using Tg276 mice.^[34]^ Because Fc-SPINK1 is large enough to prevent renal clearance, pharmacokinetics is expected to be dominated by trafficking through cells with various Fc receptors. The Tg276 mice express human versions of FcRn and the various Fc-gamma receptors, so the pharmacokinetics of Fc-SPINK1 (in which the Fc and SPINK1 elements are both based on human proteins) will be more accurately represented. The concentration of Fc-SPINK1 in blood plasma was followed over time (Figure 6). Analysis of the data using a two-compartment model indicates a beta-phase half-life of 21.51 hours, an area under the curve (AUC) of 3190 (µg·h)/ml or 132.9 (µg·days)/ml for a 100 µg dose, and a clearance rate of 0.75 ml/day.

## Discussion

Pancreatitis is an extremely painful and sometimes fatal inflammation of the pancreas in which excess trypsin activation plays a major role. To treat pancreatitis, we designed an Fc-SPINK1 fusion protein that would inhibit extracellular trypsin, and also have a long enough plasma half-life to be practical as a therapeutic protein. SPINK1 is attractive because it is a natural human protein, unlike bovine pancreatic trypsin inhibitor (BPTI/Aprotinin), which is only expressed in ruminants^[35]^ and might therefore be immunogenic. In addition, SPINK1 has a high specificity for trypsin and a much lower affinity than BPTI for other serine proteases. We placed SPINK1 at the C-terminus of the Fc region because Fc fusion proteins are generally more highly expressed in this configuration.^[23]^ We expressed this protein in the yeast *Pichia pastoris*, purified it, and showed that it inhibits trypsin *in vitro* (Figure 1).

Fc-SPINK1 reduced the indications of pancreatitis in the cerulein-induced mouse model of this disease. (Figure 2-4, Supplementary Figure 1). Cerulein, a toxic peptide from the Australian green tree frog with a C-terminus similar to cholecystokinin, hyperstimulates the pancreas and, after 7 injections leads to a mild pancreatitis that resolves after about 24 hours.^[27][30]^ When C57Bl6N mice were treated with cerulein, 18 hours after the first injection the pancreas showed signs of pancreatitis: dead acinar cells, infiltrating CD11b^+^ leukocytes, and spaces between cells that may represent edema or loss of junctional integrity. When mice received 5 mg of Fc-SPINK1 after the third cerulein injection, cell death and microscopic edema was reduced (Figures 2A-N). Blood levels of pancreatic amylase were slightly increased in cerulein-treated mice but reduced back to baseline in mice receiving Fc-SPINK1 as well (Figure 2O). These effects were observed at the higher intravenous doses of a dose-escalation experiment in which we administered 0, 1, 2.5 or 5 mgs/mouse of Fc-SPINK1 to cerulein-treated C57Bl6N mice (Figure 3). In addition, when mice of the C57Bl6J strain, which develop a more severe cerulein-induced pancreatitis than C57Bl6N mice, were treated with Fc-SPINK1 prior to the cerulein injections, a reduction in cell death and leukocyte infiltration was observed (Figure 4). Taken together these results indicate that Fc-SPINK1 can reduce the intensity of pancreatitis in the cerulein-treated mouse model.

One dramatic effect of pancreatitis in mouse models is the separation of cells in H&E-stained sections of the pancreas of cerulein-treated mice. Previously, researchers scored this by visual inspection (blinded to sample identity), which is tedious and may miss subtle effects. As part of this work, we developed an automated scoring system in which the spaces between cells in H&E stained sections are identified, abstracted as lines, and the branchpoints in the resulting pattern is quantitated (Figure 5). This approach may work in part because non-pathological spaces between cells, corresponding to ducts and spaces between pancreatic lobes, do not have extensive branching patterns and will not contribute greatly to the score.

**Figure 5.**
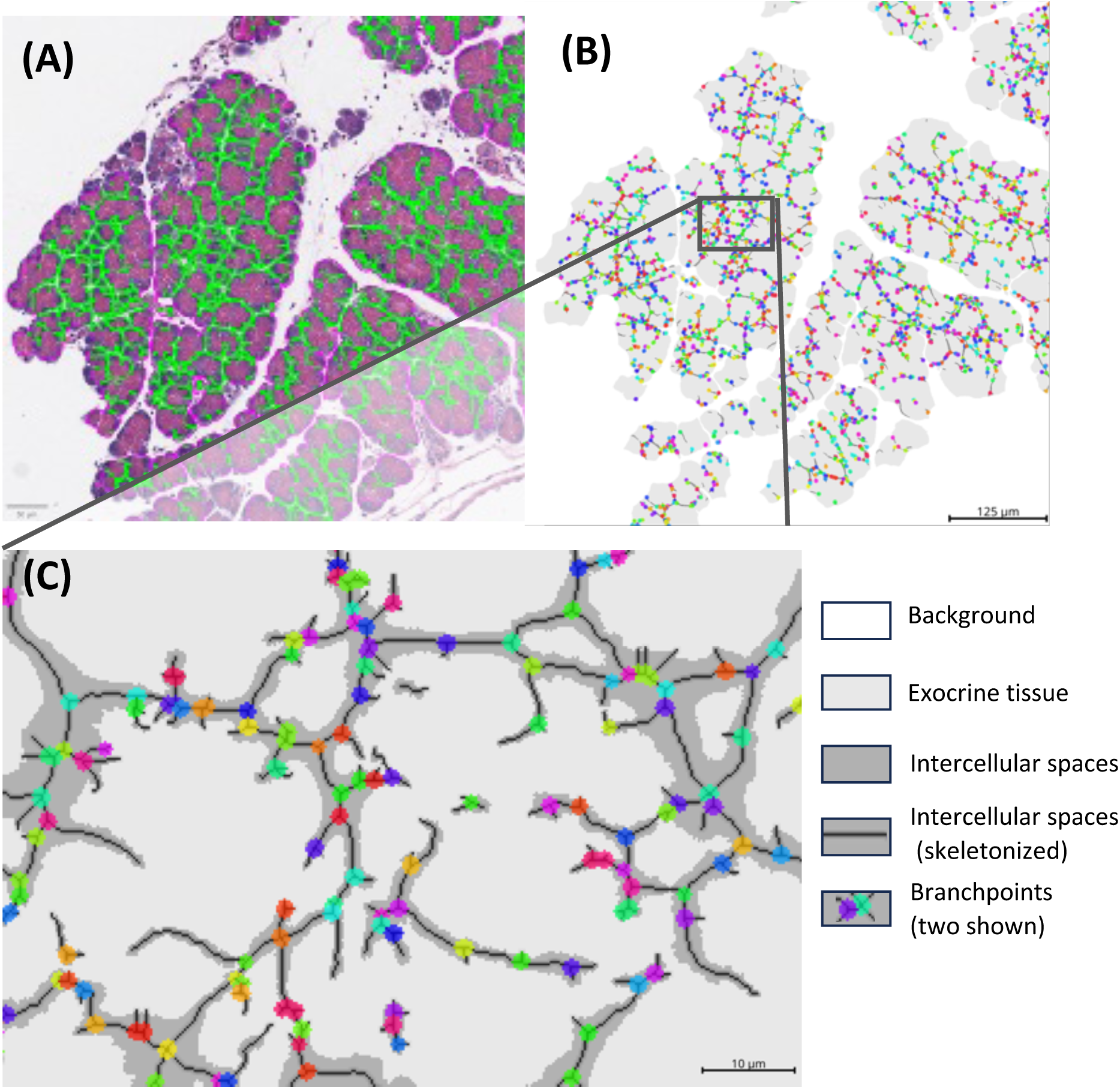
Branchpoint analysis to quantitate reduction of tissue integrity. (A) Segmented gaps between cells (B) Skeletonized gaps: the network of gaps is turned into a graph. (C) Counting the branch point of this graph serves as a proxy for how separated the cells are.

The Fc-SPINK1 protein has a long plasma half-life (Figure 6). When the data are interpreted with a two-compartment model, the elimination half-life is about 21 hours, while interpretation with a more physiological 3 compartment model indicates an elimination phase half-life of about 4 days (Supplementary Figure 4). These results suggest that the Fc-SPINK1 molecule tested here would be adequate for prophylactic administration prior to the ERCP procedure, but improvement of the plasma half-life would be ideal for chronic pancreatitis patients who may need life-long treatment.

**Figure 6:**
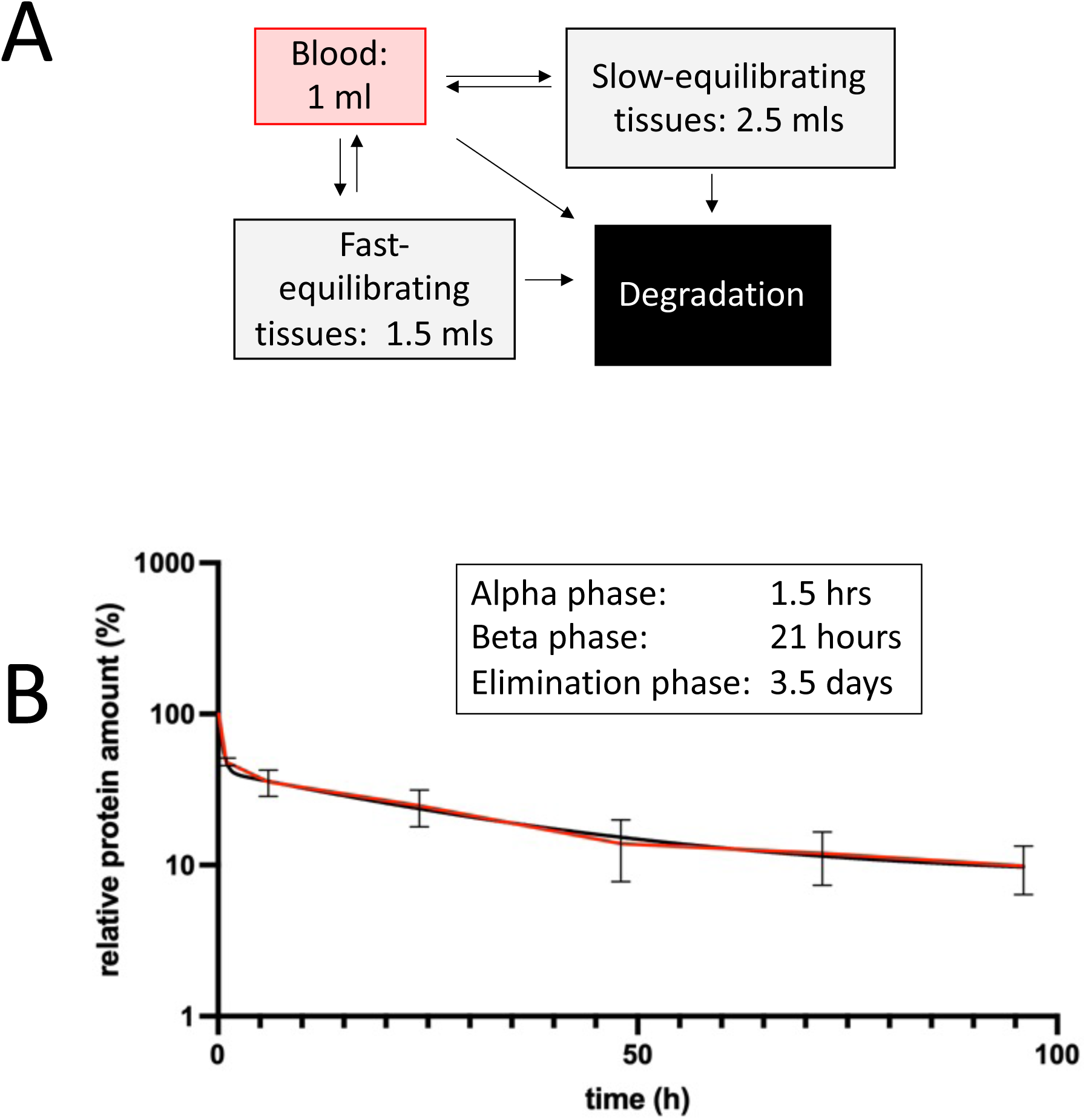
Pharmacokinetics (and distribution) of Fc-SPINK1. (A) Compartment model for estimating PK parameters. (B) Concentration of Fc-SPINK1 in the blood after IV injection of 100 mcg/mouse (N=4). The inset shows estimated pharmacokinetic parameters. The black curve is a two-parameter fit of the data. The brown lines are point-by-point straight line connections between averages of each time point.

The initiation of pancreatitis is thought to occur when overstimulation of exocrine secretion leads to fusion of acinar secretory vesicles with the wrong membrane, such as the basolateral cell surface or a lysosome or endosome. In particular, upon cerulein treatment, fusion of a trypsinogen-containing vesicle with a vesicle containing cathepsin B leads to trypsin activation within 30 minutes of treatment.^[30]^ A cathepsin B knockout mouse does not show this early activation, but still shows pancreatitis phenotypes after multiple cerulein injections. Geisz et al.^[30]^ hypothesize that trypsin autoactivation also occurs in the interstitial space and, through cleavage of protease-activated receptors, may initiate pancreatitis. The fact that Fc-SPINK1 reduces the severity of pancreatitis in the cerulein mouse model supports the idea that interstitial autoactivation plays a role in this model, and that cerulein-induced pancreatitis is not solely driven by intracellular trypsin activation.

As an injected drug, Fc-SPINK1 has advantages compared to small molecule trypsin inhibitors. Camostat is a serine protease inhibitor that is approved in Japan for treatment of pancreatitis.^[36]^ Layer et al. found that when healthy human volunteers were given oral camostat, trypsin activity was inhibited throughout the small intestine, but also increased levels of pancreatis amylase and lipase in the intestine.^[37]^ These authors inferred that this occurred through increased pancreatic secretion, and the results were evidence for a feedback loop between protease levels in the intestine and pancreatic activity. These observations suggest that any orally delivered trypsin inhibitor might be problematic for treatment of pancreatitis, since there will always be more drug in the intestine than in the pancreas, and during periods between doses when drug levels fall, pancreatic secretion could be stimulated.

A high dose of Fc-SPINK1 is required to successfully treat the cerulein-dosed mouse. This is likely because mammals produce a large amount of trypsin, and to be effective, Fc-SPINK1 needs to be in molar excess relative to active extracellular trypsin. As a point of comparison, the human body secretes between 10 and 100 mgs of trypsin per day into the small intestine.^[38–39]^ The pancreas weighs about 80 grams – slightly more than 0.1% of body weight – so only a small fraction of systemically administered Fc-SPINK1 will distribute into the pancreas. The fraction of trypsin that is activated and redirected into the accessible extracellular space during pancreatitis is unknown but may be large. With allometric scaling, a dose of 5 milligrams in a 25-gram mouse corresponds to a dose of about 1-2 grams in a human. Administration of such a large amount of protein is less challenging in humans than in mice because humans are amenable to IV infusion.

Patients undergoing an Endoscopic retrograde cholangiopancreatography (ERCP) procedure have about a 10% chance of experiencing post-procedure pancreatitis, which usually resolves but can be debilitating or fatal. Typically, before the procedure, a patient is given an IV infusion of 250 mls to 1 liter of fluid to make the ducts more accessible. Fc-SPINK1 could be administered prophylactically in this infusion to reduce the chances of developing pancreatitis. The infusion volume would allow for a large dose of Fc-SPINK1 without requiring an exotic formulation for a high protein concentration.^[40]^ The plasma half-life of Fc-SPINK1 is in an appropriate range, since post-ERCP pancreatitis occurs within 24 hours of the procedure. Taken together, our results indicate that Fc-SPINK1 is a promising candidate for treatment and prevention of pancreatitis.

## Methods

### DNA construction

Geneious Prime version 2020.1.2 was used to organize sequence data, design, and validate cloning strategies, maps, and primers. Each construct made during this research project was sequence-validated by sanger sequencing provided by GENEWIZ (South Plainfield, NJ, USA). All plasmids used in this study are based on the expression vector pPICZαA (Thermo Fischer Scientific, V19520). This plasmid allows methanol-inducible, secreted expression in *Pichia pastoris*. Plasmids were constructed by standard techniques. Fc and SPINK1 sequences were codon-optimized for expression in *Pichia pastoris*.

### Protein expression in *P. pastoris*

Expression and purification of the Fc-SPINK1 and Fc-BPTI fusion proteins was carried out using standard techniques, which are detailed in Supplementary Information. In brief, the methanol-inducible *P. pastoris* system was used to express secreted Fc fusion proteins, which were concentrated from supernatant and purified using standard Protein A chromatography.

### Protein characterization: Trypsin inhibition activity

Trypsin-inhibitor fusion proteins were tested in vitro for trypsin binding capacity. Therefore, trypsin inhibitor proteins and trypsin were diluted to a concentration of 50 µM in PBS. Trypsin was mixed with the respective inhibitor protein in ratios from 1:1 to 1:0.0625, including a positive and negative control for trypsin activity, and incubated at room temperature for 5 min.

Trypsin activity in each sample was measured using the Trypsin Activity Colorimetric Assay Kit (Abcam, ab102523). Here, trypsin activity was measured with a substrate that is cleaved by trypsin to generate p-nitroaniline (*p*-NA) which is detected at λ = 405 nm.

In brief, reactions were diluted 1:50 with PBS. Samples and controls were mixed with 50 µl reaction mix and incubated at 25°C for 60 min. Absorption at 405 nm was measured and normalized to the trypsin positive control without trypsin inhibitor

### Caerulein-induced acute pancreatitis model

Pancreatitis was induced by intraperitoneal (IP) administration of the peptide caerulein according to Lampel and Kern.^[28]^ Caerulein (50 µg/kg dissolved in 50 µl endotoxin-free PBS - 1 µg for a mouse with 20 g) was injected once per hour to a total of 7 injections. Mice were closely monitored throughout the injection period. Illness was observed within 1 – 2 hours after the first injections.

### Administration of trypsin inhibitor proteins

Trypsin inhibitor fusion proteins were administered in endotoxin-free PBS either intraperitoneal (IP) or intravenously (IV) by tail vein injection. Maximum volumes for IP and IV injections were 200 µl and 100 µl, respectively.

### Measurement of serum α-amylase activity

Serum activity of α -amylase, a digestive enzyme made by the pancreas and a clinical biomarker for pancreatitis was measured using the Amylase Activity Colorimetric Assay Kit (abcam, ab102523) according to manufacturer’s instructions.

### Histopathology: Hematoxylin and Eosin (H&E) staining and immunohistochemistry (IHC)

Immediately after necropsy, pancreas tissue was fixed in 10% Neutral Buffered Formalin for 24 hours before embedding, cutting, and staining. Slide scanning for all samples and IHC staining for CD11b was performed by Histowiz (Brooklyn, NY, USA).

### Pharmacokinetics

Pharmacokinetics of trypsin inhibitor fusion proteins were studied by IV administration of the respective protein. Tail vein blood was collected 5 min, 1 h, 3 h, 6 h, 12 h, 24 h, 72 h and 96 h post-drug injection and stored as EDTA-plasma at −20°C until analysis. ELISAs based on human SPINK1 capture and human Fc detection were used to quantitate fusion protein in serum, using standard procedures described in Supplementary Information.

### Animal welfare

All mouse experiments were carried out under protocol IS00002593, approved by the IACUC of Harvard Medical School.

## Acknowledgments

We thank Harvard Medical School for a Q-FASTR grant and the Wyss Institute for Biologically Inspired Engineering for validation project support. We also thank Andrea Geisz for extensive discussions on pancreatitis mechanisms and techniques.

## Author contributions

CW, DH, LBa and LBu were responsible for the analysis and interpretation of the data and for the preparation of this manuscript. DH, LBa, LBu, JAB, TD, PH, DN, RS, TS were responsible for protein expression, and/or performing and planning the experiments. MT and SN were responsible for imaging algorithm development, analysis, interpretation, preparation of the manuscript KR conceptualized the Fc-SPINK1 fusion protein and constructed the first versions. ARG, AV and MSV were responsible for verifying the accuracy and performance of the animal studies. PAS was responsible for funding and study supervision, and assisted with preparation of the manuscript.

## Supplementary Information

### Protein engineering and expression

To express Fc-SPINK1, DNA encoding the amino acid sequence in Figure 1 was designed with codon optimization for *Pichia pastoris* and inserted into the pPICZalpha expression vector (Invitrogen/Life Technologies) such that the EPK … coding sequence followed the … EKREAEA coding sequence in the vector, which corresponds to the cleavage site that removes the alpha-factor leader peptide that promotes secretion from yeast.

Competent cells of *P. pastoris* strain NRRL-Y-11430 were transformed using the Pichia EasyComp™ Transformation Kit (Thermo Fisher Scientific, K173001) according to manufacturer’s instructions. For secreted protein expression, 50 ml of BMGY were inoculated with the desired *P. pastoris* strain. The culture was incubated over night at 30°C, 250 rpm. The next morning, the culture was diluted to 250 ml with BMGY and incubated at 30°C, 250 rpm until they reached an OD_600_ of ∼ 5. Afterward, cells were diluted to a starting OD_600_ of 1.0 in BMMY.

The required culture volume was transferred to 50 ml conical tubes or centrifuge flasks and centrifuged at 3.000 x g for 5 minutes before resuspending in 1000 ml BMMY and transferring to a 4 l shaking flask. The culture was incubated at 30°C, 250 rpm for 48 – 96 h. Afterward, the suspension was transferred in 400 ml centrifuge bottles to separate the cells from the protein-containing supernatant by centrifuging at 3.000 x g for 15 min. The supernatant was sterile-filtered, flash-frozen and stored at −80°C or directly used for concentration.

### Protein concentration for purification

Supernatants from protein expression were concentrated before to purification by ultrafiltration using membranes with 10 kDa molecular weight cut-off at 4°C. Therefore, Centricon Plus-70 spin concentrators (Merck, UFC701008) were used according to manufacturer’s instructions to reduce the liquid volume by factor 50 - 100. The concentrates were supplemented with Halt Protease Inhibitor Cocktail (Thermo Fischer Scientific, 78429) and stored at 4°C for up to 3 days. Concentrates were flash-frozen and stored at −80°C for longer storage periods.

### Protein A affinity chromatography

IgG Fc fragment containing proteins were purified using Pierce Protein A Plus Agarose (Thermo Fisher Scientific, 22812) according to manufacturer’s instructions. Before purification, concentrated supernatants from were adjusted to pH 7.5 using concentrated solutions of NaOH or NH_4_OH. Precipitates during neutralization were removed by centrifugation at 3.000 x g for 3 min.

The clarified sample was diluted 1:1 IgG1 Binding Buffer (Thermo Fischer Scientific, 21011) before adding 1 ml Protein A Plus Agarose per 20 mg IgG Fc fragment containing protein. After incubation for 1 hour at 4°C shaking overhead, the agarose resin was collected by centrifugation for 3 min at 800 x g and loaded into a column. Alternatively, the diluted sample was directly applied to a pre-packed and equilibrated column. The column was washed with 15 column volumes Binding Buffer before eluting the target protein with IgG1 Elution Buffer (Thermo Fischer Scientific, 21004) in 1 – 4 ml fractions. The collection tubes were prefilled with 100 µl 1 M TRIS pH 8.0 per 1 ml of eluate to adjust eluted fractions to physiologic pH immediately. Elution was monitored by measuring the absorbance at 280 nm. Samples from all purification steps were subsequently analyzed by SDS-PAGE and Coomassie staining. Protein containing fractions were pooled after SDS-PAGE analysis and stored at 4°C for further use or flash-frozen and stored −80°C.

### Dialysis

Slide-A-Lyzer G2 Dialysis Cassettes (Thermo Fisher Scientific, A52972) were used for buffer exchange after protein purification according to manufacturer’s instructions. Before dialysis, large volume samples were concentrated to fit the dialysis device using spin concentrators (Centricon Plus-70, Merck, UFC701008) with a 10 kDa molecular weight cut-off. Dialysis was performed at 4°C against the desired buffer for 2 h before replacing the dialysis buffer for a second dialysis step overnight. Dialysis buffer was used at a total of at least 300 times the sample volume throughout the dialysis procedure.

### Measurement of protein concentration

Protein concentration was quantified either by BCA assay or photometric determination. The NanoDrop 2000c (Thermo Fisher Scientific, ND2000CLAPTOP) was used to analyze the absorption at 280 nm. Protein concentration was calculated according to Lambert-Beer law with the instrument’s software. The protein’s extinction coefficient required for calculation was obtained from ExPASy webtool (Swiss institute for bioinformatics). Alternatively, protein concentration was measured using the Pierce BCA Protein Assay Kit (Thermo Fisher Scientific, 23225) according to the manufacturer’s instructions in a 96-well plate.

### Endotoxin assay

Prior to in vivo application of purified proteins, the endotoxin concentration was measured using the Pierce LAL Chromogenic Endotoxin Quantitation Kit (Thermo Fisher Scientific, A39553) according to manufacturer’s instructions. The maximum acceptable endotoxin concentration for mouse studies was 0.1 endotoxin units (EU) per drug dose.

### SDS-PAGE

SDS-PAGE was used to separate proteins by molecular weight for analysis by Coomassie staining and Western Blotting. In brief, samples were diluted to an appropriate concentration and mixed with Novex Tris-Glycine SDS Sample Buffer (2x) (Thermo Fischer Scientific, LC2676) and NuPAGE Sample Reducing Agent (10x) (Thermo Fischer Scientific, NP0004) before boiling at 95°C for 3 min. Boiled samples were stored at −20°C or directly loaded into a Novex 4-20% Tris-Glycine Mini Gel (Thermo Fischer Scientific, XP04200BOX). Electrophoresis was performed at 225 V for 40-50 min in 1x Novex Tris-Glycine SDS Running Buffer (Thermo Fischer Scientific, LC2675).

### Coomassie staining

After separation of proteins by SDS-PAGE, gels were stained with Coomassie blue to visualize the protein bands. Therefore, gels were stained with SimplyBlue SafeStain (Thermo Fischer Scientific, LC6065) according to the manufacturer’s provided microwave protocol. In brief, after electrophoresis, gels were placed in 100 ml of ultrapure water and microwaved for 1 min or until the solution almost boiled. Gels were agitated on an orbital shaker for 1 min and the water discarded. This procedure was repeated 2 additional times. After the last wash, 20 ml SimplyBlue SafeStain were added, and the gel was microwaved for 45 s to 1 min or until the solution almost boiled. Gels were agitated on an orbital shaker for 5 min before discarding the stain. 100 ml ultrapure water was added to wash the gel for 10 min on a shaker. Afterward, water was replaced by 20% NaCl and the gel incubated for at least 5 min before imaging.

### ELISAs

Fusion proteins in a subset of samples were checked for overall integrity by SDS-PAGE and Western blots using antibodies targeting either Myc-tag or the respective protease inhibitor. In this enzyme-linked immunosorbent assay (ELISA) protocol designed for the detection of SPINK1, the process unfolds systematically for optimal results.

To initiate the procedure, the capture antibody (anti-SPINK1; abcam, ab207302) is diluted at a ratio of 1:1000 in coating buffer (8.4 g/l NaHCO_3_, 3.56 g/l Na_2_CO_3_ in H_2_O). Subsequently, 100 µl of the diluted antibody is added to each well, followed by an overnight incubation at 4°C.

Moving on to the blocking phase, the plate is brought to room temperature, and each well undergoes five washes with wash buffer (0.5 ml/L Tween-20 in PBS).

200 µL of blocking buffer (5% BSA in PBS) is then added to each well and incubated for 1 hour at room temperature. Another round of five washes with wash buffer follows.

Standards and samples are diluted in sample buffer (1% BSA in PBS). Then 100 µl of each diluted standard or sample is added to the appropriate wells and the plate is incubated for 2 hours at room temperature. This is followed by another five washes with wash buffer.

For the detection antibody step, biotin-labeled detection antibody (goat anti-human IgG Fc (biotin); abcam, ab97223) is diluted to 200 ng/mL in sample buffer. 100 µl of the diluted antibody is added to each well and the plate is incubated for 1 hour at room temperature. This step is followed by five washes with wash buffer.

Streptavidin-Horseradish Peroxidase (SA-HRP) is added by diluting the conjugate 1:15,000 in sample buffer. 100 µl of the diluted conjugate is added to each well and the plate is incubated for 30 minutes at room temperature. Simultaneously, the TMB substrate is prepared by placing the required amount in a conical tube, wrapping it in aluminum foil, and allowing it to reach room temperature in the dark for 30 minutes. The plate is then washed 5-8 times with PBS/Tween.

To add the substrate (TMB), 100 µl of equilibrated TMB is added to each well using a multichannel pipette. After waiting 1-5 minutes or until the desired dynamic range is reached, 100 µl of 1 M sulfuric acid is added to stop the reaction.

The final step is to read the optical density (OD) for each well using a microplate reader set at 450 nm.

### Image analysis

#### Code availability

https://doi.org/10.5281/zenodo.14894990

#### Edema tutorial

https://github.com/HMS-IAC/quantify_edema

### Explanation of batch identifiers

Three imaging experiments were performed

Experiment 1: Identifiers 11571 and 11564 refer to a batch referenced in Figure 2. Experiment 2: Identifiers 14628 and 14550 refer to the dose response experiments. Experiment 3: The identifier 25868 refers to a batch of mice of strain C57Bl6J.

### Segmentation

Segmentation is the process of assigning pixels to a class, such as endocrine tissue, exocrine tissue, necrosis, and edema. Segmentation was performed using the histopathology software QuPath: Tissue examples are annotated manually. These annotations were used as ground truth to train a machine-learning based pixel classifier. For reproducibility, pixel classifier settings are saved as a .json file and applied in .groovy scripts, which can be run from QuPath (see code availability). Such obtained segments were manually corrected where specified.

### Segmentation of exocrine and endocrine tissue

Unless specified otherwise, pixel classifiers were trained on a composite image from samples of batches 11571, 14550, and 25868. The pixel classifier endo_exo_bin_20240521.json was trained using QuPath and used to segment endocrine and exocrine tissue using the script fine_grained_segmentation.groovy. For exocrine tissue, minor manual corrections were made by removing wrongly segmented (false positive) areas. Endocrine tissue was manually corrected by both removing false positive segmentations and correcting false negative segmentations. Next, the segmentation was processed to achieve a smoother outline to accelerate downstream computations: The polygon representing the segmentation was simplified by removing vertices (corners), it was expanded by 15 microns to fill holes, then shrunk by 15 microns. If exocrine and endocrine tissue overlapped after these operations, the overlapping area was assigned to endocrine. (coarse_grained_segmentation.groovy).

### Segmentation of Edema

Edema is brighter (higher grayvalue) than surrounding tissue. Hence, bright background may mistakenly be segmented as edema. Hence, bright background pixels were segmented based on the thresholder background.json and subtracted from the class exocrine. Exocrine tissue is shrunk by 1 um to remove additional background. A pixel classifier edema_20241006.json was trained on a composite image of a randomly selected 200 x 200 um region from each image using training_annotation.groovy and used to segment edema within exocrine tissue using refine_exocrine.groovy.

### Segmentation of Necrosis

For 14550 and 25868, exocrine tissue segmentation from the previous step (Edema) was used. The pixel classifier necrosis_ignorebackground.json was trained on a composite image of 200 x 200 um regions selected from every other image, including images from batch 11571. These training regions were randomly selected and, if necessary, manually shifted to a region containing necrotic cells. The pixel classifier necrosis_ignorebackground.json was applied to the bounding box of exocrine using segment_necrosis.groovy. Necrosis segmentation that overlapped with endocrine was assigned to endocrine instead using mergenecrosis.groovy. False positive segmentations were manually removed. The area (um^2^) of exocrine tissue and necrotic tissue is exported to CSV using export_necrosis.groovy.

### Segmentation of Necrosis in 11571

To achieve a more precise segmentation of necrotic tissue in 11571, pixel classifiers were trained on 11571 specifically. The previous exocrine annotation was removed and endocrine tissue was subtracted from the exocrine boundingbox. Within the exocrine bounding box, two 500 x 500 um regions were randomly selected for each image in 11571 and shifted to a region with necrotic cells, where necessary. These regions were combined into a training image (training_annotation.groovy) to train the pixel classifier exocrine11571.json. Within the exocrine boundingbox, new exocrine tissue was classified using pixel classifier exocrine11571.json. Background from pixel classifier background.json was subtracted (new_exocrine.groovy). Next, pixelclassifier necrosis2.json was specifically trained on 11571 and applied to segment necrosis within the exocrine region (segment_necrosis.groovy). False positive segmentations were manually removed. The area (um^2^) of exocrine tissue and necrotic tissue is exported to CSV using export_necrosis.groovy.

### Segmentation of CD11B

**14628**: The pixel classifier tissue_ignorebg.json was trained to segment tissue areas in batch 14628, immune_cells_ignorebg.json was trained to segment immune cells within the segmented tissue of 14628. (segment_exocrine.groovy). The relative area of immune cells to tissue was computed.

**11564**: The pixel classifier exocrine.json was trained to segment tissue areas in batch 11564, immune_cells.json was trained to segment immune cells within the segmented tissue of 14628 (segmentation11564.groovy). The relative area of immune cells to tissue was computed.

### Exporting Edema

The edema mask and exocrine mask were exported as .png files. Batch 14550 and 25868 were exported at full resolution. Batch 11571 was exported at 1.5x downsampling (export_edema_experimental.groovy) due to computational limitations. Images were split into tiles (tile_image.py) to enable further processing. For each tile, the edema mask was skeletonized, and the branch points of the resulting skeleton were counted. The ratio of branchpoints to exocrine area, based on the exocrine mask, was computed (edema_to_graph_downsampled.py).

### Exporting and quantifying necrosis

Per image, the ratio of the necrotic area to exocrine area was calculated (analyse_necrosis.py).

**Supplementary Figure 1.**
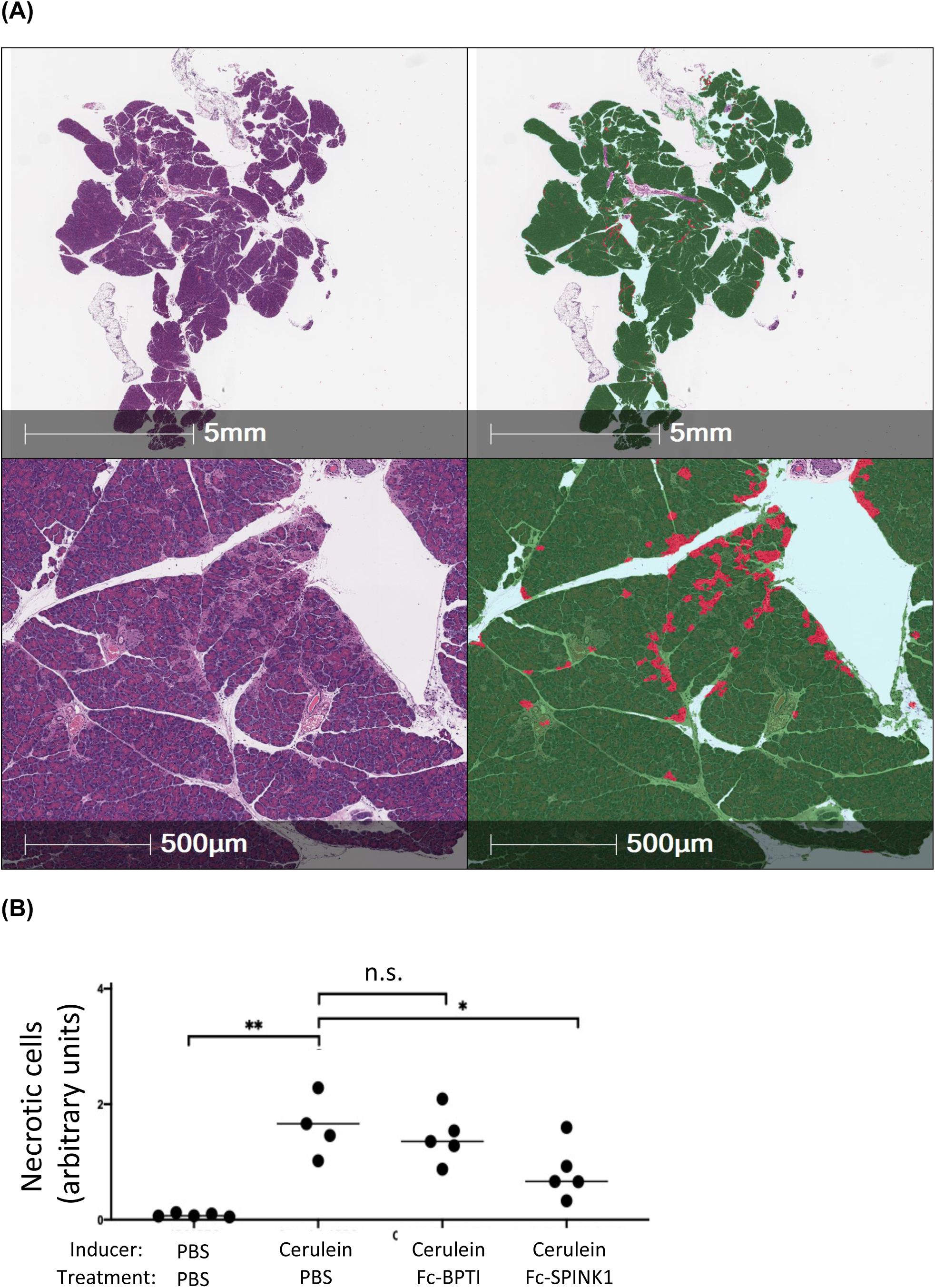
**(A)** Quantification of necrotic cells in hematoxylin/eosin-stained pancreas sections from mice using an alternative automated image analysis method. These images were generated by Histowiz, a commercial histology provider, using their proprietary image analysis pipeline. **(B)** The same data as analyzed for Figure 2N by our image analysis software was also analyzed by the Histowiz method (n.s. – not signficant; * p<0.05; ** p<0.01: one-tailed Student‘s T-test). Results from the Histowiz analysis are consistent with those generated by the image analysis approaches described above, supporting the validity of our open-source approach

**Supplementary figure 2:**
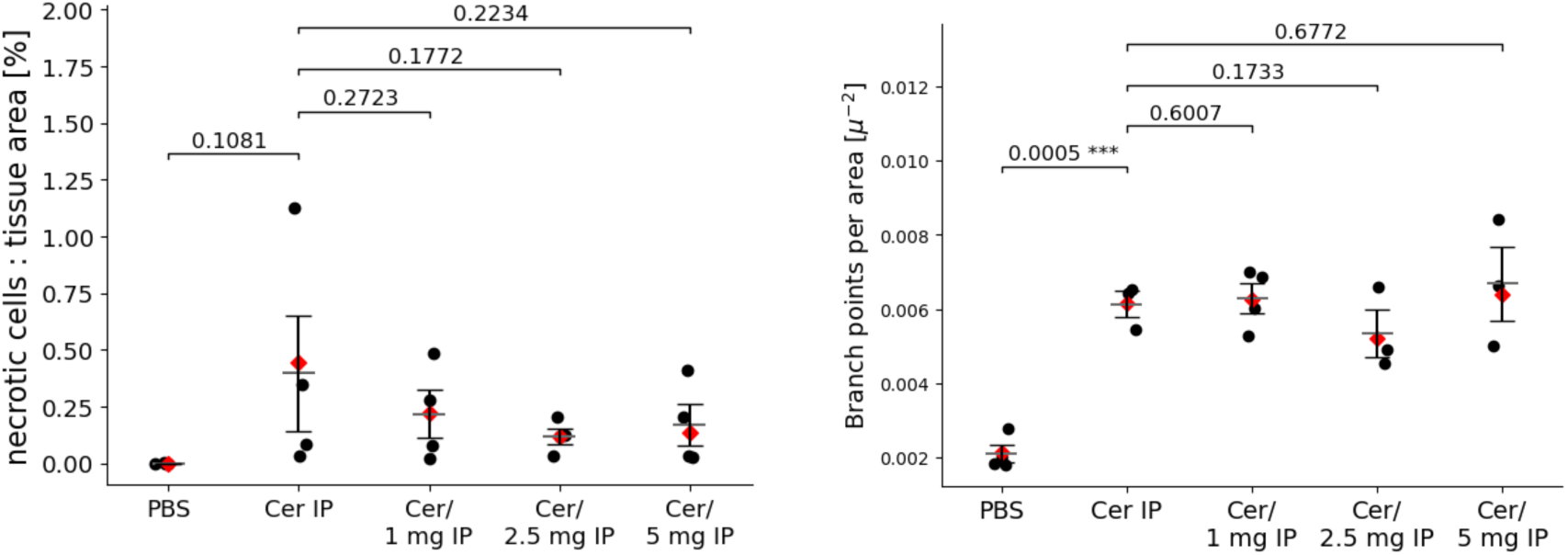
IP dosing in dose escalation. A and B: Horizontal bars indicate mean ± standard error of mean (SEM). Red diamonds indicate mean weighted by tissue-area. P-values: single-tailed Welch’s t-test. **A:** Results of semi-automated necrotic cell staining. Pancreases of PBS-treated mice show no necrotic cells, while pancreases of cerulein treated mice showed a increased level of cell necrosis. **B:** Analysis of tissue integrity.

**Supplementary Figure 3:**
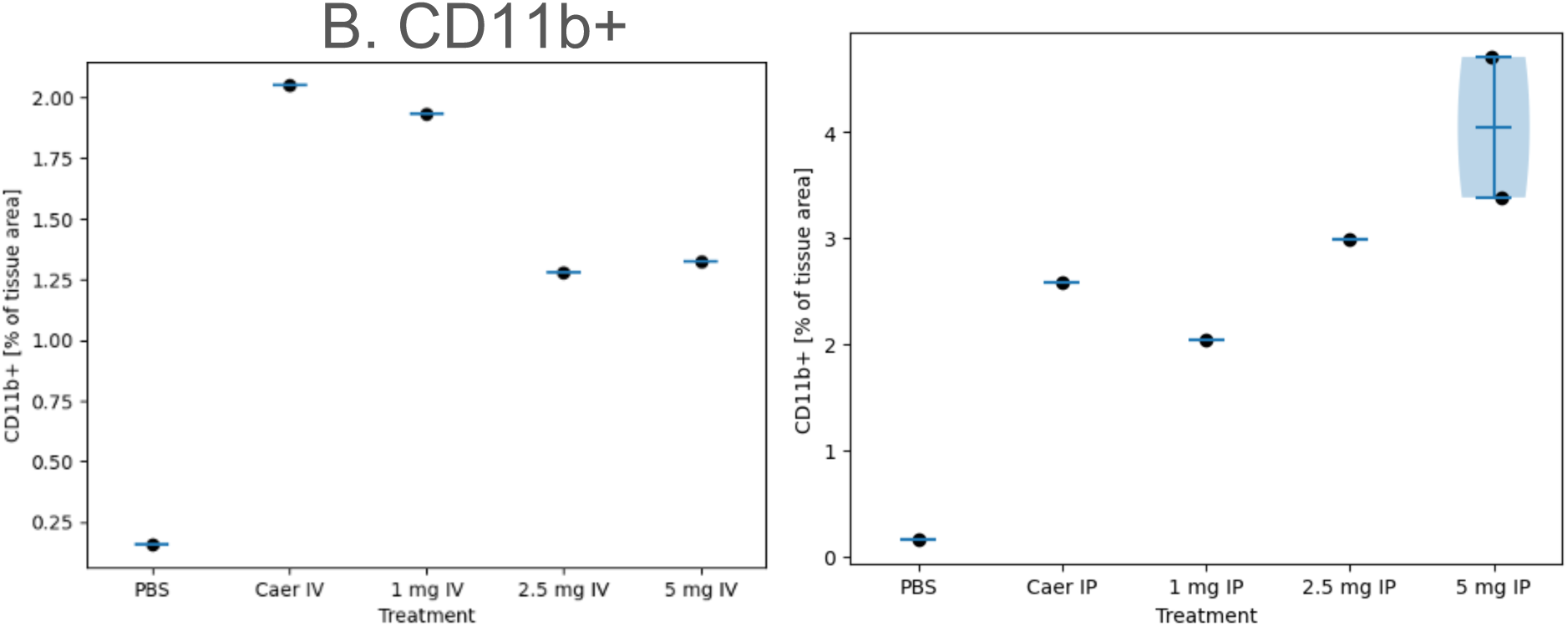
Image analysis of CD11b+ leukocyte infiltration in the dose response experiment. A subset of cross-sections from the experiments in Figure 3 and Supplementary Figure 2, respectively were stained with anti-CD11b+ antibodies imaged using HRP immunohistochemistry, and analyzed using the automated image analysis methods described above.

### Note on the pattern of Fc-SPINK1 inhibition of trypsin

We tested every batch of Fc-SPINK1 for trypsin inhibition and generally found a step-function increase in inhibition, without signs of partial inhibition at molar ratios around 0.5:1 of SPINK1:trypsin. This is observed in Figure 1F. Buchholz et al. found that there are actually two binding sites for SPINK1 on trypsin. It may be that since Fc-SPINK1 is dimeric for SPINK1, it can form a higher-order complex with a high Hill coefficient of binding.

Reference: Buchholz, I., et al. The impact of physiological stress conditions on protein structure and trypsin inhibition of serine protease inhibitor Kazal type 1 (SPINK1) and its N34S variant. BBA – Proteins and Proteomics. 1868,1-8 (2020).

**Supplementary Figure 4:**
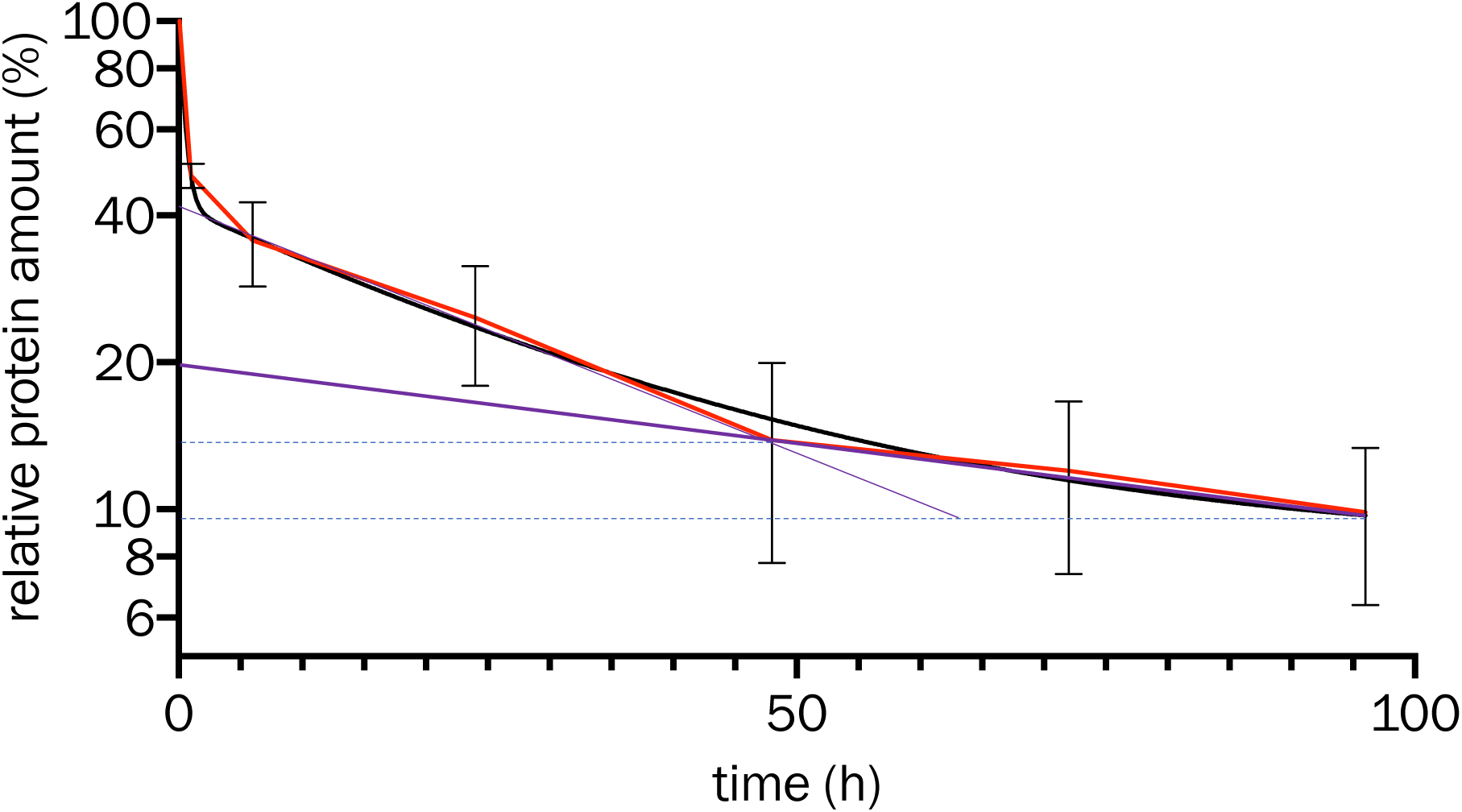
Pharmacokinetics of Fc-SPINK1, analyzed by a three-parameter fit. Garzone and Atkinson^[1]^ have argued that the pharmacokinetics of protein drugs that do not bind to or penetrate cells should be modeled with a three-parameter fit, corresponding to the plasma compartment, a ‘fast’ equilibrating compartment corresponding to splanchic tissues such as the liver, which have fenestrated capillaries, and a ‘slow’ compartment, corresponding to the rest of the tissues. Here we take the 48-hour timepoint to represent the point at which distribution into the fast compartment is complete. The last three timepoints suggest a terminal plasma half-life of about four days, since the relative protein amount decreases from about 14% to about 9.8% in two days.

1. Garzone, P.D. & Atkinson A.J. In search of physiologically based distribution volume estimates for macromolecules. Clin. Pharmacol. Therapeutics 92,419-421 (2012).

